# Determinants of variation in epidemiological parameters for African Swine Fever virus

**DOI:** 10.1101/2025.07.21.665936

**Authors:** Marian Talbert, Abigail Feuka, Ryan S. Miller, Kim Pepin

## Abstract

African Swine Fever (ASF) is a highly infectious reportable disease of swine that can have devastating consequences for pork producers and consumers. ASF virus can persist in either domestic or wild swine through several epidemiological cycles. This has led to a wide variety of epidemiological traits, making it challenging to plan effective surveillance and mitigation strategies. Here, we compile and analyze data from experimental infections of ASF virus variants from genotype II that have been reported in peer-reviewed publications. We provide descriptive summaries of the distributions of epidemiological parameters such as incubation period, infectious period, mortality rate, and R_0_ and develop predictive models that explain the variation in these quantities based on detection assays and other experimental design features. Our results provide a comprehensive perspective of estimates of epidemiological parameters for ASFV, allowing for increased transparency in accounting for parameter uncertainty in ASF preparedness modeling. Our meta-analysis also provides insight on knowledge gaps and study design issues that could be addressed by future experimentation.

## Introduction

African swine fever virus (ASFV) causes the most economically harmful disease of swine globally. The continued global spread of ASFV has significantly reshaped global protein markets, created supply gaps for human protein, and altered animal feed protein markets (Carriquiry et al., 2019; Mason-D’Croz et al., 2020; Schmidhuber et al., 2020). This has resulted in significant economic losses in countries affected with reported costs and/or losses resulting from ASFV outbreaks ranging from $796,000 to $116 billion (USD 2024) (Bech-Nielsen et al., 1993 and Fasina et al., 2012). For this reason, many countries are investing substantial effort into controlling ASFV where it is endemic or preventing its introduction in countries that are currently ASFV-free (Animal Health Australia 2022, United States Department of Agriculture 2023, European Commission 2024). Implementing these plans effectively relies on strong knowledge of ASFV epidemiological characteristics – including incubations period, infectious period, mortality rate, and the basic reproductive number of ASFV (R_0_) in different host population contexts.

ASFV is a large, double-stranded DNA virus in the *Asfarviridae* family. It exclusively infects *Suidae* (MacLachlan et al., 2017). This virus evolved in eastern and southern Africa as part of a sylvatic cycle involving warthogs (*Phacochoerus africanus*) and soft-bodies ticks of the genus *Ornithodoros* (Penrith and Vosloo, 2009; Bakkes et al., 2018). There are four known epidemiological cycles through which ASFV can transmit and persist in swine, including one that involves wild boar and the environment (Chenais et al., 2018). It is the only known DNA virus causing hemorrhagic fever transmitted by an arthropod, and can result in high mortality in susceptible populations of wild and domestic swine (Blome et al., 2013).

Genotype II viruses are the cause of the ongoing global pandemic that started in 2007 (Zhang et al., 2023). Genotype II viruses were introduced into Georgia in 2007 from sub-Saharan Africa and have been identified in many ASFV outbreaks occurring outside Africa since 2007 (Malogolovkin et al., 2015, Shen et al., 2022, Zhang et al., 2023). Genotype II spread from Georgia to neighboring countries (Armenia, Azerbaijan, Russia, and Belarus) where a highly virulent ASFV strain infected domestic and wild swine (Karger et al., 2019). At least nine countries in the European Union (Estonia, Lithuania, Latvia, Poland, the Czech Republic, Bulgaria, Belgium, Romania, and Hungary) have had persistent spread despite significant efforts to control transmission in both wild and domestic swine hosts (WOAH, 2024). Many of these countries are now considered to have endemic transmission (Jiang et al., 2022) with wild boar playing an important role in local spread and maintenance (Sauter-Louis et al., 2021). Genotype II subsequently spread from Europe to Asia, Oceania, and the Americas. ASFV was identified in China in 2018 where it has been difficult to eliminate, and the Western Hemisphere for the first time since 1980 with continuing outbreaks in the Dominican Republic and Haiti since 2021 (Wang et al., 2018, Spinard et al., 2023; WOAH, 2024).

Twenty-four genotypes of ASFV have been identified with genotype II viruses, highlighting substantial genetic variation within the genotype group. More recently variation in morbidity and mortality has been observed within genotype II viruses (Gallardo et al., 2018). ASFV isolated from wild boar in Estonia, Lithuania, and Russia can cause a longer course of infection and asymptomatic infections with recovery (Vlasova et al., 2014; Gallardo et al., 2017; Nurmoja et al., 2017). For example, surveillance of apparently healthy wild boar has found antibody-positive wild boar demonstrating that some proportion of animals survive infection (Woźniakowski et al., 2016). Also, experimental infections using AFSV strains from Estonia (Es15/WB-Valga-6 and Es15/WB-Tartu-14) found moderate virulence with surviving animals demonstrating a lack of viremia and clinical signs (Gallardo et al., 2018). Similarly, the strain of ASFV (ASFV-DR21) isolated from the Dominican Republic does not consistently produce an acute and fatal form of disease (Ramirez-Medina et al., 2022).

Studies investigating progress of disease and differences among virus genotypes or strains using experimental infection are valuable for characterizing epidemiological parameters such as length of incubation and infectious periods, mortality rates, or transmission to naïve contact animals. Experimental studies are advantageous because they allow characterization of the course of infection within individuals based on a known time of infection and quantification of the level variation in epidemiological parameters (Sanchez-Cordon et al., 2019). However, inconsistencies in experimental methodologies and reporting can limit inference by making comparisons across studies challenging, which could improve weight of evidence (Blaustein et al., 2018). A lack of standardization in experimental methods, particularly for infection studies, has been identified as a limiting factor in generalizing about disease dynamics for many host-pathogen systems (Kilpatrick et al., 2010; Blaustein et al., 2018). To reconcile potential differences in study design and assays used for measuring pathogen infection time courses, sources of variation across experiments need to be accounted for to allow generalization and estimation of epidemiological parameters that govern disease dynamics.

A current challenge with experimental work on ASFV has been the lack of standardization in experimental design, assays, and reporting of study results. Here, we develop statistical models of published epidemiological parameters from experimental infection studies of ASFV that account for differences among studies to generalize about four principle epidemiological measures - incubation period, infectious period, recovery rate, and transmission rate. Our primary objective is to provide estimates of these parameters that account for a realistic range of potential variation to equip disease managers with an appropriate perspective of the uncertainty in ASFV epidemiological dynamics. This is important because epidemiological models are often used to guide preparedness or control using a narrow range of epidemiological parameters – for example Halasa et al., 2016 and Lee et al., 2021. Our approach summarizes the known variation in ASFV epidemiological parameters in a predictive model that can be used in risk assessment and preparedness applications to obtain more realistic transparency of uncertainty. We also provide a discussion of the current limitations of available experimental studies for ASFV.

## Materials and Methods

### Data collection

We conducted a systematic search of the peer-reviewed literature to identify articles estimating parameters associated with incubation period, infectious period, recovery rate, and two correlates of transmission rate (daily transmission rate, β rate and basic reproductive number, R_0_) for ASFV in wild boar or domestic swine. Our literature search is outlined in Fig. 1. We restricted our search to papers published after January 1^st^, 1980, and before January 1^st^, 2020. The search was implemented using three databases (PubMed, Scopus, and Web of Science). We extracted a list of articles that contained either the phrase “African Swine Fever” or “ASF” in the title or keywords and contained one of the words incubation, infection, infectious, experimental inoculation, recovery, transmission, contact rate, dynamics, or epidemiology. We then merged the resulting article lists based on DOI and removed duplicates. We further restricted to papers reporting results on ASF virus genotype II because it is the most widely distributed ASF virus variant and are the viruses currently involved in the ongoing global pandemic. We then restricted to experimental studies on swine and observational studies of outbreaks that did not genetically modify the ASF virus strain, did not vaccinate, and did not remove infected pigs to slow transmission in transmission experiments. To identify papers potentially missed by our initial search, we also reviewed the citations of papers identified as relevant to our study, papers citing articles in our study, as well as those in review articles identified in the initial literature search.

**Fig. 1.**
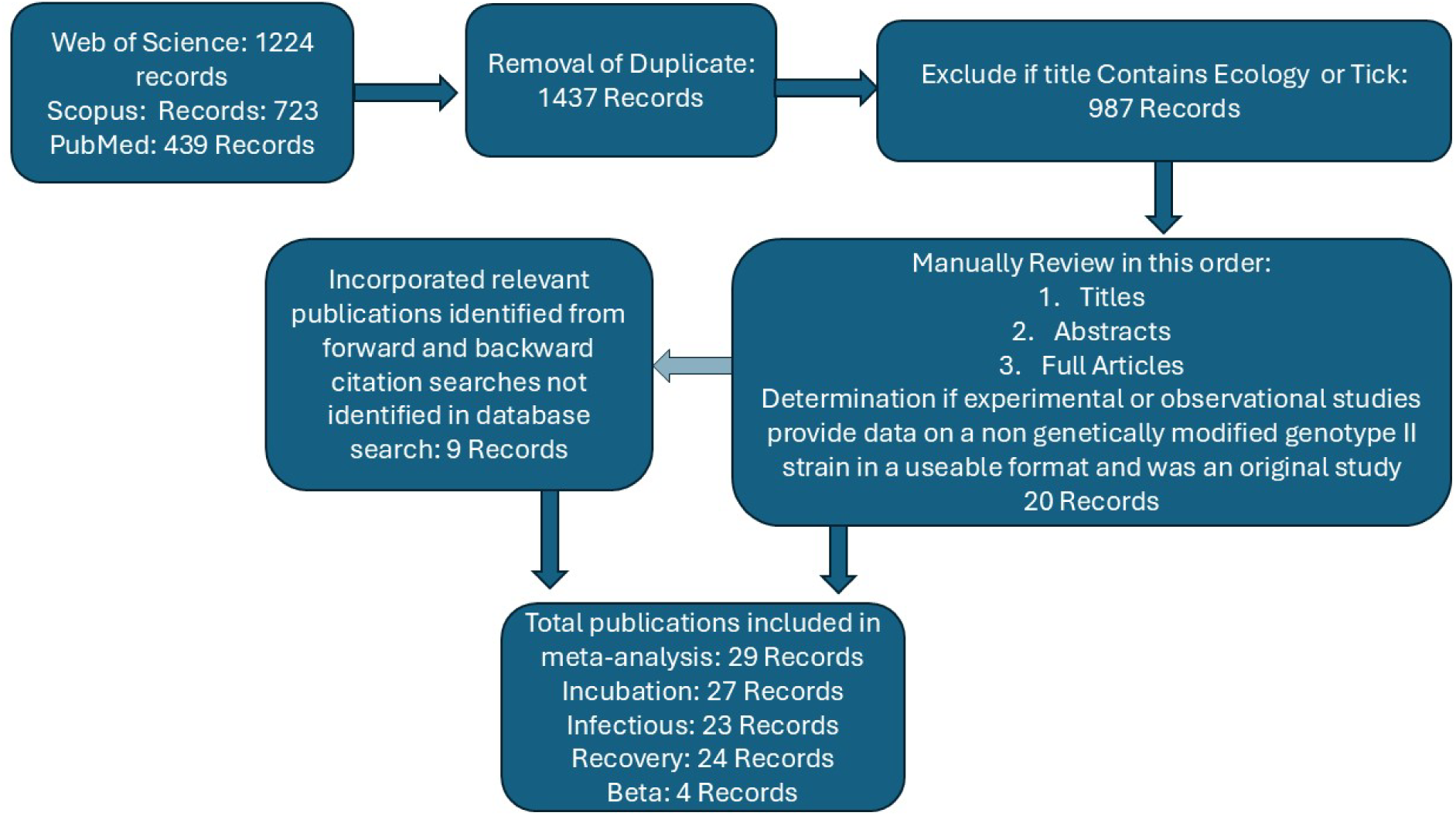
Flowchart outlining the article selection process carried out to identify data sources.

### Data description and processing

Data reported for all measurements (e.g. animal temperature, viral levels in excretions/blood, observed clinical signs, etc.), animal identifier, and study identifier were recorded. We also recorded characteristics describing age, sex, and species of animals, virus characteristics (strain), as well as aspects of study methodology and diagnostic assays used (Table 1).

**Table 1.**
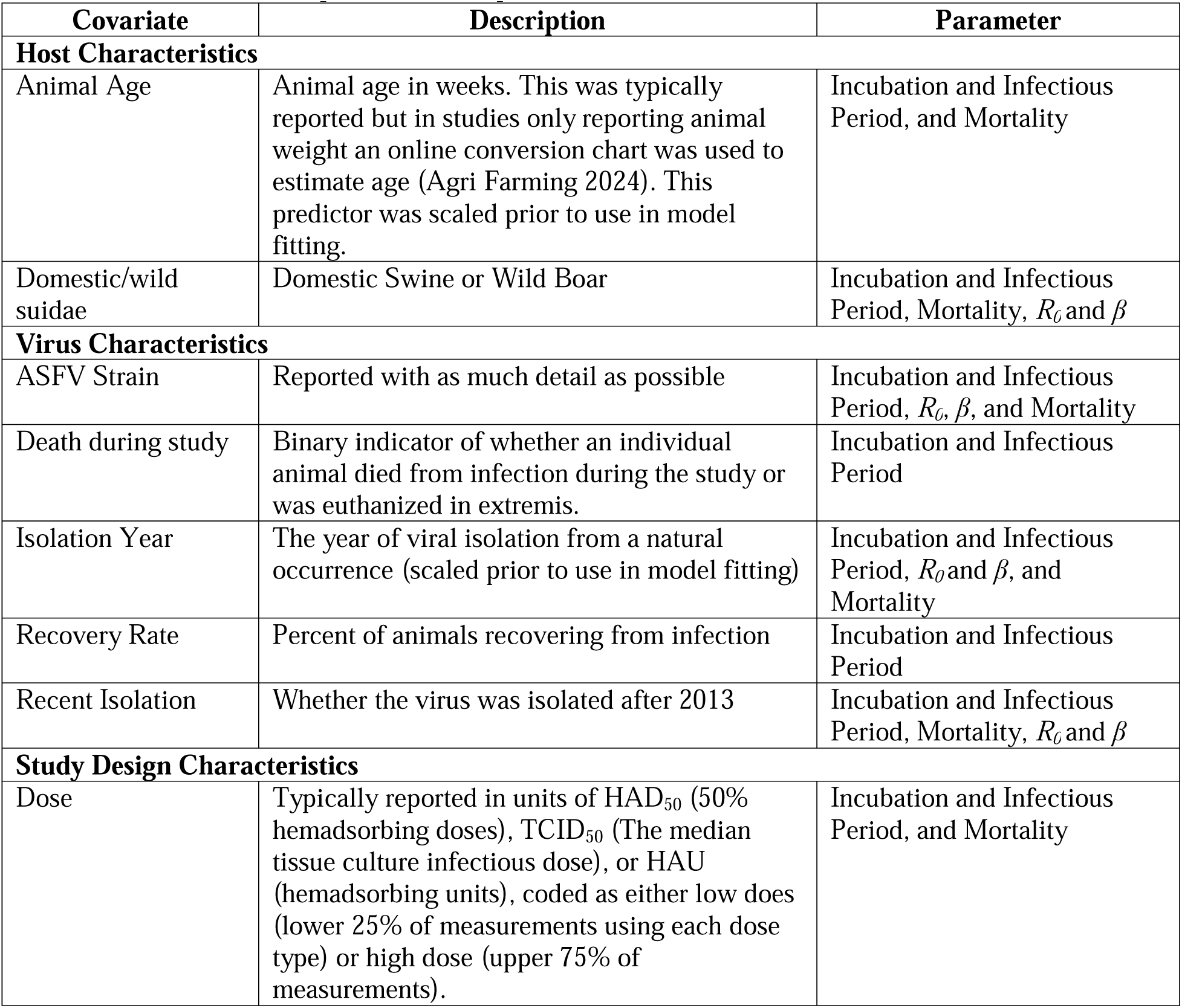
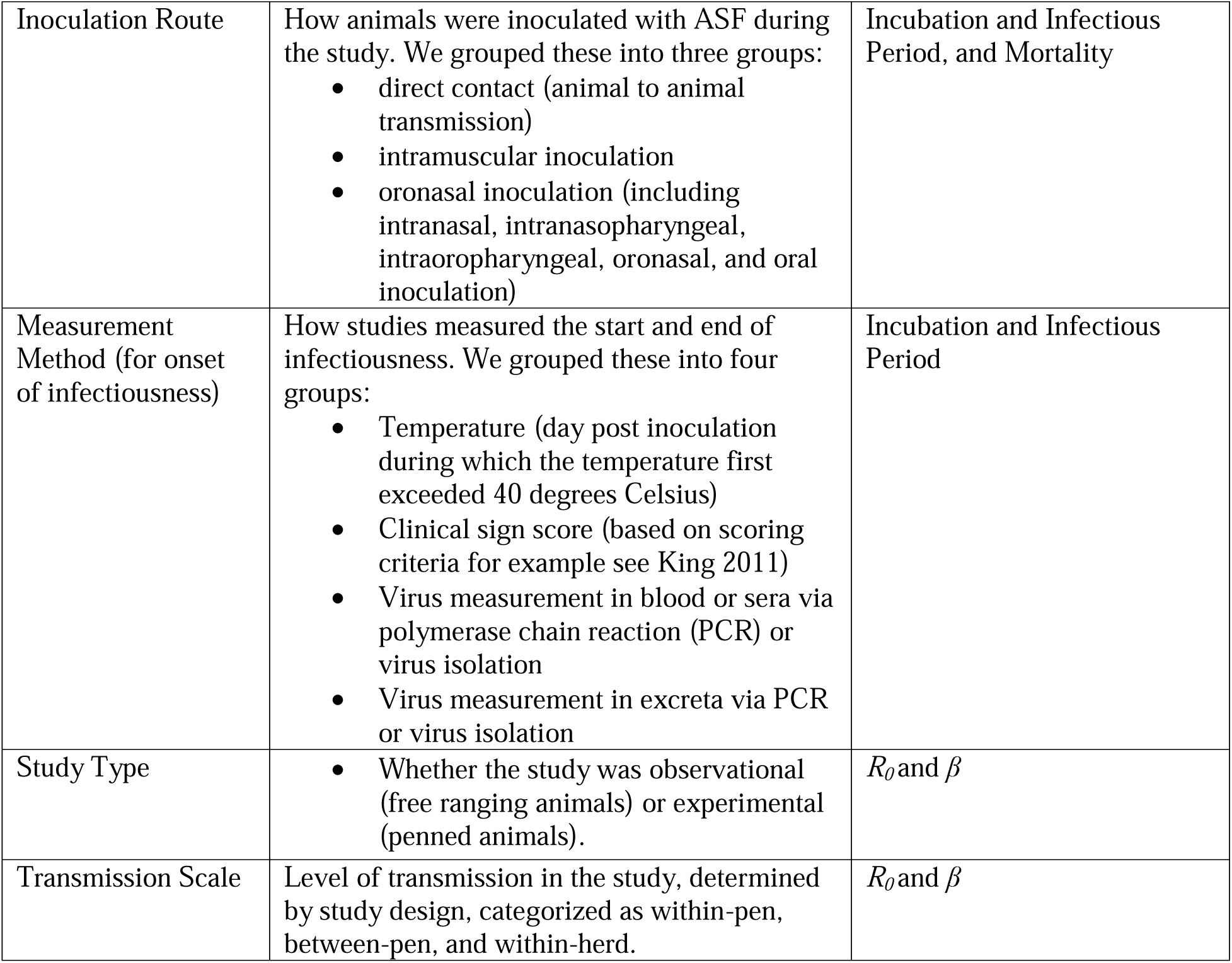
Covariate descriptions and the parameter models each covariate was used to estimate.

**Table 2.** Data availability for fitting transmission rate.

| **Inoculation Route** | **How Determined** | **Unique Articles** |
| --- | --- | --- |
| Between Pen | Model to fit experimental data | Guinat et al. (2016), Hu et al. (2017), Nielsen et al. (2017) |
| Within Herd | Model to fit observational data | Guinat et al. (2018), Le et al. (2023), Mai et al. (2022b) |
| Within Pen | Model to fit experimental data | Guinat et al. (2016), Hu et al. (2017), Nielsen et al. (2017) |

To capture individual variation more accurately, we used individual-level data from studies when it was available or could be inferred from plots or supplementary material. When studies only reported group-level distributions, we simulated individual observations using a program evaluation and review techniques (PERT) distribution parameterized by the minimum, maximum, and mode reported in the study.

Available individual-level data was often panel censored, in that samples were not collected daily and the testing interval often changed during the study. When exact days of onset and termination of infectious period were not available, these were imputed by stochastic simulation from a uniform distribution to randomly select a time of onset of infectiousness between the sample collection times. In studies where the first sample was collected later than the potential onset of disease (i.e., day three or later after inoculation), we simulated the onset of infectiousness using a beta distribution putting higher weight toward the three-day time period (drawing from Beta(shape1=4, shape2=1) and multiplying this by the date of first detection minus the left panel limit, and adding date of first detection). For the most common scenario in which the animal was detected on the first day of sample collection (day three) and the logical lower limit for onset of infectiousness is zero, this function assigns three days as the onset 53% of the time, day two as the onset 41% of the time, and day 1 as the onset 6% of the time. When virus was detected intermittently, we used the last day of detection to indicate the end of the infectious period and day of euthanasia was used when animals were euthanized in extremis or due to study conclusion.

Incubation period was defined as the days between inoculation and onset of viral shedding, detection in blood, clinical signs score, or temperature depending on how incubation period was recorded by the specific study. For incubation period models, observations associated with the low-dose oronasal and direct contact inoculation route were removed, as well as any records above the 97.5% quantile of onset for intramuscular inoculation. This decision was made because not all animals become infected from the initial inoculation, but often through secondary contact with their pen-mates and in these cases the date of inoculation is unknown (e.g., Pietschman et al., 2015).

Infectious period was defined as the time between the end of the incubation period and either death or last detection of clinical signs, temperature or viral shedding depending on how it was measured. Models were fit separately for animals surviving infection and those dying either from the infection or euthanized in extremis because the latter represent a truncated view of infectious period.

### Data analysis

#### 1. Predictor Selection

Exploratory data analysis was carried out to provide empirical distribution summaries, identify potential relationships, determine how parameters should be modeled, and develop groupings for categorical predictors. A visual inspection was performed to assess relationships between pairs of categorical predictors and relationships between continuous and categorical predictors. Any continuous covariates with Pearson correlation coefficients greater than 0.7 were not used in the same model.

#### 2. Model Fitting

Linear Mixed Models (LMMs) were fit to the log-transformed incubation and infectious period respectively using the “lmer” function from the “lme4” package (Bates et al., 2015) with nested random effects for studies and subjects within study. We conducted backwards stepwise selection using the “lmerTest” package (Kuznetsova et al., 2017) and used an F-test for marginal fixed effects terms to select the most parsimonious models. The significance of random effects for both study and animal were assessed using likelihood ratio tests using the “rand” function from “lmerTest”. Models were evaluated by consideration of residual patterns and based on marginal and conditional R^2^ values calculated using the “r.squared” LMM function in the “MuMIn” package (Bartoń 2023).

In addition to the LMMs, we fit probability density functions to infectious period and incubation period observations to quantify individual animal-level variation grouped by factors found to be most influential in modeling. Distributions were fit using the “fitdistrplus” package in R (Delignette-Muller et al., 2015). We fit gamma distributions because the parameters were continuous and bounded above zero. Distribution fits were assessed using the Kolmogorov-Smirnov test (Massey 1951) with the “stats” package in R (R Core Team 2024).

Recovery rate was modeled as a generalized linear mixed model using the “glmer” function from the “lme4” package using the bounded optimization by quadratic approximation (“bobyqa” optimizer option, Bates 2015) with a Bernoulli response indicating whether individual animals recovered. The most parsimonious set of predictors was selected via AICc using the “dredge” function in the “MuMIn” package. Because animal age was typically held constant within each study, we used this as a covariate rather than using random effects for each study. Models were evaluated using Area Under the receiver operating Curve (AUC) values with the “pROC” package (Robin et al., 2011) and conditional and marginal R^2^ values calculated using the “r.squared” function in the “MuMIn” package.

Variability in assumed or estimated infectious period can be highly influential in estimates of R_0_ (Schulz et al., 2019). To avoid issues with this source of variability, we analyzed the transmission rate, β, rather than R_0_. The log transformed transmission rate, log(β), was modeled for within pen, between pen, and within herd spread using the meta-analysis methodology provided by the “Metafor” package (Viechtbauer 2010) for linear mixed effects models. The log transformation was used to ensure the estimates would be bounded on the positive interval.

Because the number of animals involved was drastically different for experimental versus observational studies, the number of herds rather than the number of animals was used as the sample size for observational studies and the number of replications was used as the sample size for experimental studies. Model selection was carried out using the “dredge” function in the “MuMIn” R package.

## Results

### Literature search

Our search of PubMed, Scopus, and Web of Science resulted in 1,437 articles associated with AFS after duplicates were removed. Limiting papers to those meeting our inclusion criteria resulted in 987 papers which were manually reviewed to determine applicability. Manual review resulted in a final set of papers included 27 for incubation period, 23 for infectious period, 24 for recovery rate, and 4 for transmission rate. Studies included in our analysis are listed in Supplemental Table S1.

### Incubation Period

The intercept term for incubation period, exp(β) (95% CI) = 5.47 (4.83, 6.18), captures the mean animal age of 10.4 weeks inoculated with a low dose with measurements taken via temperature and clinical signs. Significant predictors for incubation period included host age exp(β) (95% CI) = 1.03 (1.01, 1.05), measurement via blood exp(β) (95% CI) = 0.88 (0.81, 0.96), via excretions exp(β) (95% CI) = 1.30 (1.19, 1.41), and high dose exp(β) (95% CI) 0.70 (0.62, 0.80). The marginal pseudo R^2^ for the fixed effects component of the best model was 0.28.

Incorporating random effects for study and animal within study increased the conditional pseudo R^2^ to 0.55. Fig. 2A shows the effect sizes for incubation period coefficients and detailed information on model selection and model fit parameters are provided in Supplemental Tables S2 and S3.

**Fig. 2.**
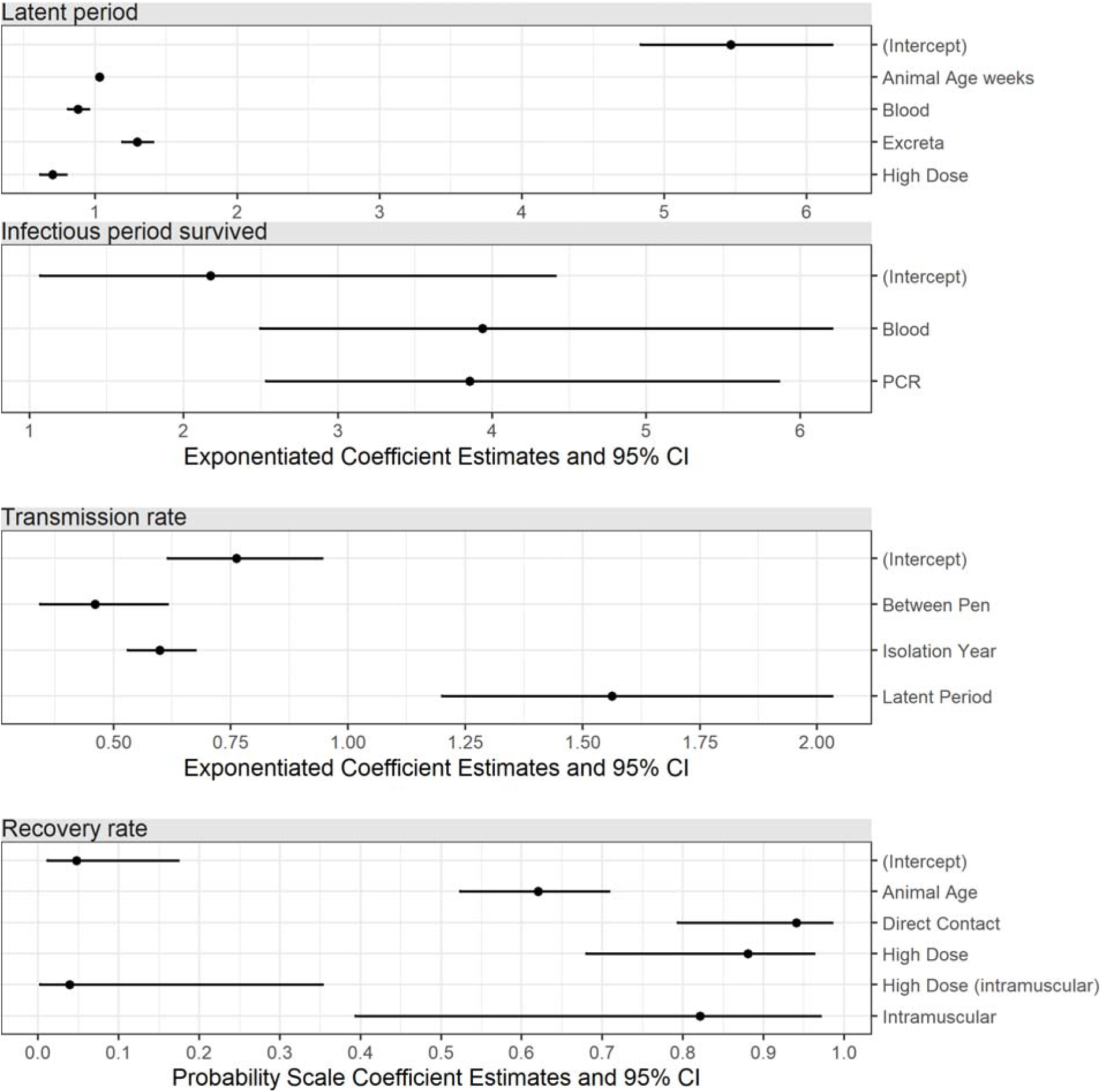
Effect size plots from top to bottom for A) incubation period, B) infectious period for animals that survived, C) transmission rate, and D) recovery rate, transformed to the raw scale.

### Infectious Period

Both the length of infectious period and the factors governing infectious period were drastically different for animals surviving versus succumbing to ASF infection (Fig. 6). For this reason, we elected to fit separate models to these two groups. For animals dying from ASF the pseudo R^2^ for the fixed effects component of the best model was 0.07, indicating that the available predictors were not able to explain much of the variation. Incorporating random effects for study and animal within study increased the proportion of variation that was explained to 0.71 indicating that random effects were the primary driver of infectious period in animals dying of ASF. Given the low R^2^ for fixed effects, we elected to not report parameter estimates for this model.

There were very few animals surviving ASF infections and even fewer for which clinical signs or temperature data were available, so modeling and distribution fitting was only possible for infectious period for animals that survived and when infectious period was measured via blood and excreta. For this group, the two significant predictors were measurement method and virus detection method. The intercept term represented infectious period for excreta measured by virus isolation exp(β) (95% CI) = 2.17 (1.07, 4.41). Infectious period was far longer when determined using blood exp(β) (95% CI) = 3.94 (2.5, 6.21), especially when measured via PCR exp(β) (95% CI) = 3.85 (2.54, 5.86). The pseudo R^2^ for the fixed effects component of the best model was 0.34 and incorporating random effects for study and animal within study increased this to 0.53. Fig. 2B shows the effect sizes for infectious period in surviving animals. Detailed information on model selection and model fit parameters are provided in Supplemental Tables S4 and S5.

### Recovery rate

There were no records in our meta-analysis for an animal experimentally inoculated and surviving with a genotype II virus strain that was isolated prior to 2013 despite approximately 40% of the data coming from this period. The pseudo R^2^ for the fixed effects and random effects components were 0.31 and 0.47 respectively. The effects plot shows inverse logit transformed estimates and confidence intervals (Fig. 2D). The intercept term, inverse logit(β) (95% CI) 0.05 (0.01, 0.17), represents recovery rate for an animal of age 10 weeks inoculated oronasally with a low dose. Recovery rate was higher for animals infected via direct contact inverse logit(β) (95% CI) 0.94 (0.79, 0.98) and increased with scaled animal age inverse logit(β) (95% CI) 0.62 (0.52, 0.71). Both high dose and intramuscular inoculation had higher recovery probabilities than the intercept inverse logit(β) (95% CI) 0.88 (0.68, 0.96) and inverse logit(β) (95% CI) 0.82 (0.39, 0.97) respectively but the combination of high dose and intramuscular inoculation was associated with much lower probability of recovery inverse logit(β) (95% CI) 0.04 (0.03, 0.35). Detailed information on strain specific mortality rate, model selection and model fit parameters are provided in Supplemental Tables S6 through S8.

### Transmission rate

Data for estimation of transmission rate, β, was sparse with very few experimental studies. Hu et al. (2017) and Nielsen et al. (2017) both estimated transmission parameters using experimental data provided by Guinat et al. (2016), and Guinat et al. (2018) estimated transmission rate using three different assumptions for length of incubation period using the same experimental data.

Because nearly all data on between and within pen transmission rate was based on a single experimental study and lacked independence, the formal meta-analysis should be interpreted cautiously. Observational data for within-herd spread was available for multiple outbreaks. Two of these studies provided estimates for multiple herds. Guinat et al. (2018) analyzed outbreak data from nine fattening pig operations in Russia with herd sizes ranging from 600 to 2,145, Mai et al. (2022b) analyzed data from seven small farms (100-299 swine) and three operations with inventories between 300 and 999 swine in Vietnam, Le et al. (2023) analyzed data from a single farm in Vietnam with 17 swine. The intercept term for transmission rate, exp(β) (95% CI) = 0.76 (0.61, 0.94) comprises transmission for within pen and within herd for the mean assumed or estimated latent period of 5.6 days for strains isolated in 2010. The most significant covariate was between-pen transmission with a much lower transmission rate than within pen or with herd exp(β) (95% CI) = 0.46 (0.34, 0.61) and longer latent periods were associated with higher transmission estimates exp(β) (95% CI) = 1.52 (1.2, 2.03). The coefficient for isolation year was exp(β) (95% CI) = 0.60 (0.53,0.67) (Fig. 2C). Information on model selection and model fit parameters is provided in Supplemental Tables S9 and S10.

### Distributions for incubation period

The distributions for incubation period were very different for each of the three inoculation routes and outliers were present in the data. Fig. 6A shows the incubation period data following removal of observations for direct contact inoculation, low dose oronasal inoculation, and observations above the 97.5% quantile of intramuscular observations in gold. Once these outlier observations were removed a single distribution, shown in blue, was fit to all remaining data the shape and rate parameters for the fits were 9.99 ± 0.72 SE and 2.12 ± 0.16 respectively leading to a distribution with a median of 4.5 days and 95% of the distribution falling between 2.2 to 8.0 days (2.5% to 97.5% quantiles).

### Distributions for infectious period

There were marked differences in the potential infectious period length between animals that survived versus animals that either died or were euthanized in extremis due to their ASFV infection (Fig. 6B through 6F). The median length of infectious period was 4 days for animals that died and 14.5 days for surviving animals. For animals that ultimately succumbed to the disease, few factors were influential in determining the length of infectious period, thus a single distribution was fit to data from all animals that died. For animals surviving, there was insufficient data to fit distributions when clinical signs or temperature were used as an indicator of infectious period length, and different distributions were fit for each combination of PCR or viral isolation (VI) and for blood or excretions, as these factors were all highly significant in model fitting (Fig. 2B). All information for distribution fits is provided in Supplemental Table S11 including shape and rate parameters with corresponding standard errors, distribution medians 2.5% quantiles and 97.5% quantiles.

## Discussion

Experimental infection studies have revealed a wide range of individual-level outcomes of ASFV infection. A better understanding of the breadth of potential epidemiological parameters that determine outbreak dynamics of ASFV is important for providing an appropriate level of transparency in risk assessments and developing effective contingency plans in outbreak response guidelines. We summarize the available science describing epidemiological parameters of ASFV at the individual animal level and identify the properties that influence them in a quantitative framework. Our analysis provides a predictive framework for extracting epidemiological parameters of ASFV with appropriate uncertainty for disease transmission models that can be used for planning outbreak response or control.

Some recent studies, such as those carried out by Ramirez-Medina et al. (2022) and Gallardo et al. (2021) provide information at the individual animal level on PCR and virus titers in both blood and various swabs or excreta, in addition to daily temperature. Both studies also followed animals over long periods of time, ameliorating some of the issues associated with panel censoring and elucidating the potential contributions of long-term carriers in disease spread.

Information of this quality is valuable for comparing virus dynamics in wild boar versus domestic pigs, as these relate host age, inoculation route, and dose, allowing an understanding of variation among individuals through the course of infection. A key challenge with meta-analysis of experimental infection data that are collected using a variety of assays is that PCR-based measures of viral infectious period are often longer than measures based on viral isolation and are likely overestimates of infectious period because PCR can detect inactive viral sequences (Pepin et al., 2023). Similarly, studies differed by their PCR methods and viral isolation procedures, which can differ in sensitivity (Gallardo et al., 2015) and leads to observational variation in estimates of epidemiological parameters in addition to variation due to the true underlying process. Our modeling approach allows this observational source of variation to be accounted for in our parameter estimates.

Another challenge with characterizing the true individual-level variation in epidemiological parameters is that most experimental studies were carried out on the Georgia 2007 strain (13 of the papers used in the analysis), yet many recent outbreaks have been attributed to strains with lower virulence (lower mortality rates). Thus, there is an information bias towards ASFV epidemiological parameters in our data set where epidemiological traits of the Georgia 2007 strain are overrepresented relative to other less virulent strains. Also, some longer-term studies showed intermittent virus detection in excreta and blood of less virulent isolates (Sun et al., 2021, Petrov et al., 2018), however, with panel censoring it was not always possible to determine the exact number of days these animals were infectious, and, in some studies, viral titers were not reported, so determining how infectious an animal might have been during a second acute phase was not possible. In the case of intermittent viral detection, we used the last day of virus detection, acknowledging that several studies question how infectious these animals truly are after the first acute period (Ståhl 2019 et al., and Wilkinson 1984). On the other hand, some studies euthanized animals at the end of the study precluding the possibility of observing a second acute phase. It is generally thought that these long-term carrier animals contribute minimally to disease dynamics (Bloome et al., 2020, Ståhl et al., 2019) but given that many recent outbreaks did not match predictions based on our understanding (Pejsak and Tarasiuk 2022) of the parameters, more research is needed in these areas.

Estimated daily transmission values for the calibration data ranged from 0.31 to 0.46 with a mean of 0.38 for between pens and 0.24 to 1.58 with a mean of 0.83 for within pens. Using our median values of infectious period for animals that die and survive infection (4 and 14.5 days, respectively), this would approximately translate to R_0_ values ranging from 0.97 to 6.34 with a mean of 3.32 within pen and 1.23 to 1.83 with a mean of 1.5 between pens for an animal that dies, and from 3.52 to 23.0 with a mean of 12.0 within pen and 4.46 to 6.62 with a mean of 5.44 between pens for an animal that survives infection for a completely susceptible population.

Comparisons to other between pen, within pen and within herd studies are shown in Fig. 4 and 5 for β and R_0_ respectively. For domestic pigs, a study in Uganda used multiple methods to estimate the basic reproductive number and obtained estimates ranging from 1.58 to 3.24 (Barongo et al., 2014). In the Dominican Republic, the between-farm reproductive ratio was estimated to range from 1 to over 15 depending on the infectious period (1 to 40 days) and day of the study, with R_0_ peaking between days 1 and 25 of the outbreak (Schambow et al., 2023).

**Fig. 3.**
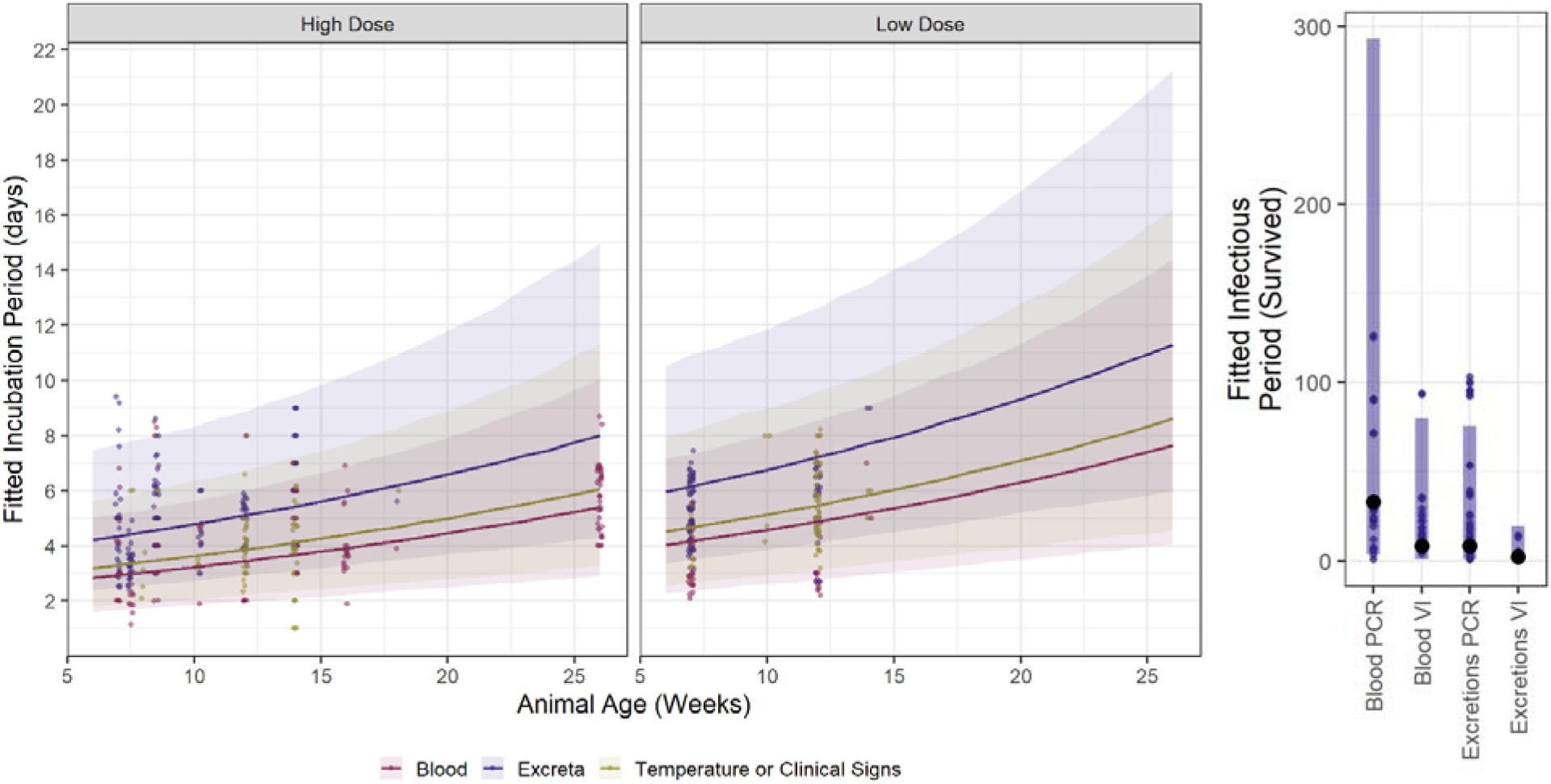
Linear mixed model fits back transformed with 95% prediction intervals showing predictors from the final model for incubation period. The data used in model fitting are included as jittered points to allow discrimination of individual data. B) Model fits for infectious period for animals surviving by measurement method and virus detection method (VI = viral isolation and PCR=polymerase chain reaction). Large circles show the fitted values, bars show the 95% prediction intervals and smaller circles show the observational data.

**Fig. 4.**
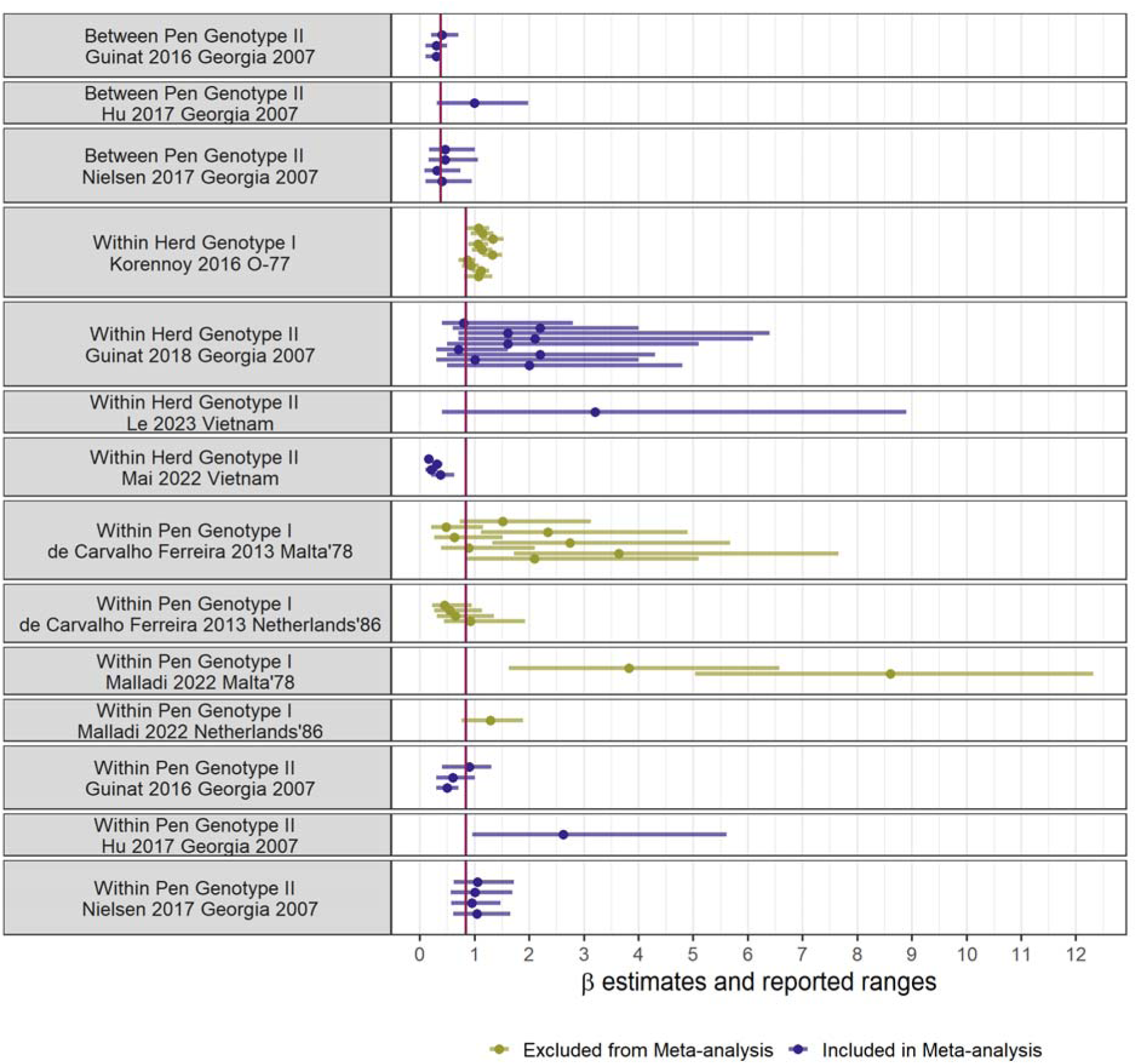
β estimates (dots) and reported ranges (lines) from previous studies. Multiple lines per reference indicate that estimation was caried out on multiple herds (Guinat et al., 2018), using multiple model assumptions, parameters, doses, and/or outbreak waves (Guinat et al., 2016, Nielsen et al., 2017, de Carvalho Ferreira et al., 2013, Korennoy et al., 2016, Malladi et al., 2022) or for different farms and pig types (Mai et al., 2022). The red vertical line shows the mean predicted value from the current study’s meta-analysis for the two groups (Between pen for the top 3 studies and within herd/ within pen for all other studies).

**Fig. 5.**
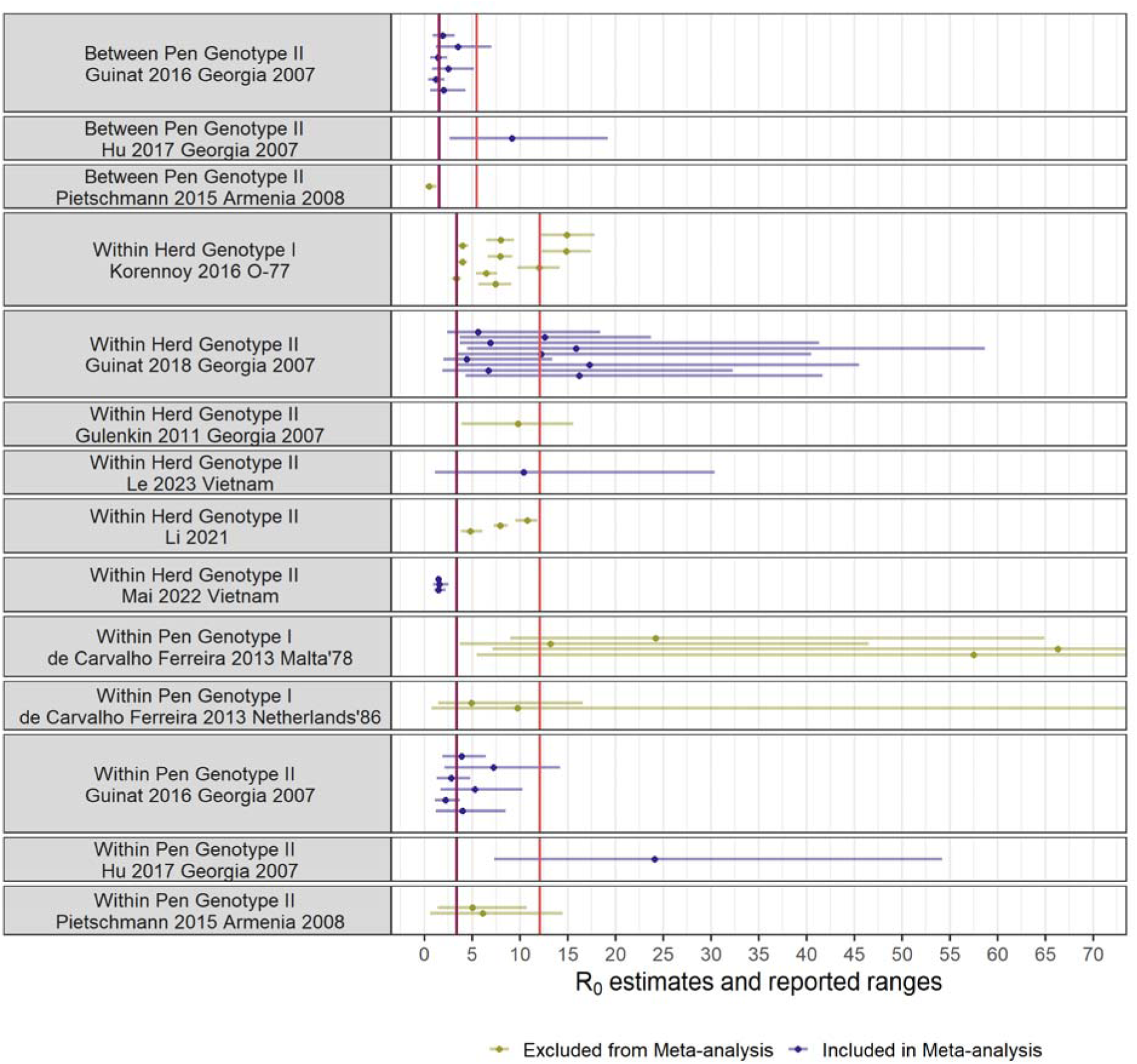
R_0_ estimates (dots) and reported ranges (lines) from previous studies. Multiple lines per reference indicate that estimation was caried out on multiple herds (Guinat et al., 2018 and Li et al., 2021), different experimental groups (Pietschmann et al., 2015), using multiple model assumptions, parameters, doses, and/or outbreak waves (Guinat 2016, Nielsen et al., 2017, de Carvalho Ferreira et al., 2013, Korennoy et al., 2016), or for different farms and pig types (Mai et al., 2022). The red vertical line shows the mean predicted value from the current study’s meta-analysis for the two groups (Between pen for the top 3 studies and within herd/ within pen for all other studies).

**Fig. 6.**
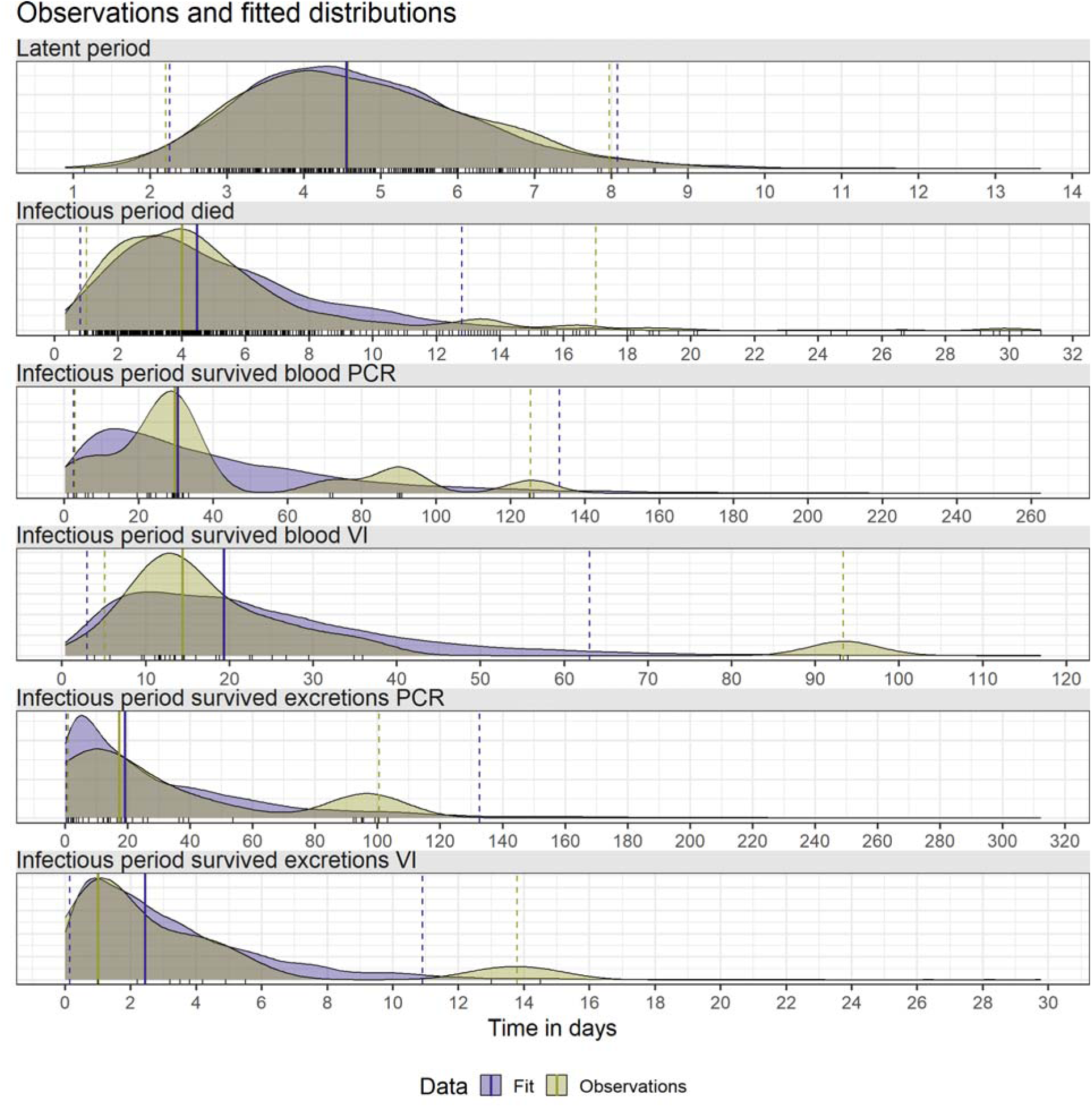
A) Distributions for incubation period of ASFV in swine by inoculation. Observations above the 97.5% quantile of the intramuscular inoculation value were removed B) infectious period of ASFV in swine for animals that died, C) infectious period for animals that survived and measured via blood with polymerase chain reaction, D) infectious period for animas that survived when measured via blood with viral isolation, E) infectious period for animals that survived when measured via excreta with polymerase chain reaction, and F) infectious period for animals that survived when measured via excreta with viral isolation. There were insufficient data to fit a distribution for surviving animals using temperature or clinical signs.

Estimates of R_0_ are lower for free-ranging wild boar that interact with fewer susceptible individuals. One study of fresh wild boar carcasses basic reproduction rate to be 1.41 and 1.66 in two locations (Loi et al., 2022). For wild pigs, study on wild boar in eastern Poland estimated the effective reproductive numbers between 1.1 and 2.5 (Pepin et al., 2020). For wild boar in the Russian Federation, transmission was estimated to be 1.33 to 3.77 (Iglesias et al., 2014). These estimates are generally within our wide range of values of approximate R_0_, which accounts for differences in isolation year and latent period. A few exceptions are the two studies based on moderately virulent genotype I strains for which a very long infectious period of between 20 and 44 days were assumed (de Carvalho Ferreira et al., 2013 and Malladi et al., 2022).

ASF transmission is complex, with multiple factors influencing when and where transmission can occur. For studies of within-herd spread, it is difficult to know how parameters may vary in other contexts because farming practices and other local factors that drive transmission dynamics (Bellini et al., 2021, Juszkiewicz et al., 2023, Ramachanderen et al., 2024) can vary dramatically. As pointed out by Cross et al. (2005), population structure is extremely important in describing the course of disease spread through a population. Thus, multiple observational studies conducted across a breadth of contexts are useful for estimating transmission rates in natural settings and understanding how different factors affect them.

## Conclusion

We provide a comprehensive analysis of epidemiological parameters of ASFV that can be used to infer ASFV transmission with appropriate uncertainty. Our analysis describes variation in epidemiological parameters from study design characteristics, highlighting how these factors affect inference. Our approach summarizes the available data in predictive models that can be leveraged to extract the best available information on parameters of interest for a given set of conditions.

## Acknowledgements

We thank Jessie Trujillo for providing virological perspective that guided the direction of this work in the early stages.

