## Supplementary Information for "Determinants of variation in epidemiological parameters for African Swine Fever virus"

### Supplement

Table S1. Studies used in the analysis and whether data was available for estimation of each parameter.

| Source | Incubation | Infectious | Recovery | $R_0$ | β | Strain(s) | DOI |
|---|---|---|---|---|---|---|---|
| Borca et al. (2017) | Yes | Yes | Yes | | | Georgia 2007 | 10.1038/srep46747 |
| Borca et al. (2020) | Yes | Yes | Yes | | | Georgia 2007 | 10.1128/JVI.02017-19 |
| Borca et al. (2021) | Yes | Yes | Yes | | | Georgia 2007 | 10.3390/v13050765 |
| Elsukova et al. (2017) | Yes | Yes | Yes | | | Odintsovo 02/14 | 10.25125/agriculture-journal-IJOEAR-OCT-2017-15 |
| Gabriel et al. (2011) | Yes | Yes | Yes | | | Caucasian | 10.3201/eid1712.110430 |
| Gallardo et al. (2017) | | Yes | Yes | | | Lithuania LT14/1490 | 10.1111/tbed.12346 |
| Gallardo et al. (2018) | Yes | | Yes | | | Es15/WB-Valga-6, Es15/WB-Tartu-14 | 10.1016/j.vetmic.2018.04.001 |
| Gallardo et al. (2021) | Yes | Yes | Yes | | | Pol16/DP/OUT21, Est16/WB/Viru8, Lv17/WB/Rie1 | 10.1111/tbed.14222 |
| Guinat et al. (2014) | Yes | Yes | Yes | | | Georgia 2007 | 10.1186/s13567-014-0093-8 |
| Guinat et al. (2016) | | | | Yes | Yes | Georgia 2007 | 10.1017/s0950268815000862 |
| Guinat et al. (2018) | | | | Yes | Yes | Georgia 2007 | 10.1111/tbed.12748 |
| Hu, Gonzales, and Gubbins (2017) | | | | Yes | | Georgia 2007 | 10.1038/s41598-017-17174-8 |
| Le et al. (2023) | | | | Yes | Yes | Vietnam 2019 | 10.1186/1471-2105-12-77 |
| Lee et al. (2021) | Yes | | Yes | | | VNUA/HY/Vietnam | 10.1186/s40813-021-00215-0 |
| Li et al. (2022) | | | | Yes | | China 2018 | 10.1111/tbed.14345 |
| Nurmoja et al. (2017) | Yes | Yes | Yes | | | Northeastern Estonia | 10.1111/tbed.12614 |
| O'Donnell, Holinka, Krug, et al. (2015) | Yes | Yes | Yes | | | Georgia 2007 | 10.1128/JVI.00969-15 |
| O'Donnell, Holinka, Gladue, et al. (2015) | Yes | Yes | Yes | | | Georgia 2007 | 10.1128/JVI.00554-15 |
| O'Donnell et al. (2017) | Yes | Yes | Yes | | | Georgia 2007 | 10.1128/JVI.01760-16 |
| Olesen et al. (2017) | Yes | Yes | Yes | | | POL/2015/Podlaskie/Lindholm | 10.1016/j.vetmic.2017.10.004 |
| Pershin et al. (2019) | Yes | Yes | Yes | | | 15 Strains originating from the Russian Federation 2013-2018 | 10.3390/vetsci6040099 |
| Pietschmann et al. (2015) | Yes | Yes | Yes | Yes | | Armenia 2008 | 10.1007/s00705-015-2430-2 |
| Pikalo et al. (2021) | Yes | Yes | Yes | Yes | | Belgium 2018 | 10.3390/ani11092602 |
| Ramirez-Medina et al. 2(022) | Yes | Yes | Yes | | | Dominican Republic strain | 10.3390/v14051090 |
| Sun et al. (2021) | Yes | Yes | Yes | | | HLJ HRB1 20 | 10.1007/s11427-021-1904-4 |
| Vlasova et al. (2015) | Yes | Yes | Yes | | | Kashino 04/13, Boguchary 06/13, | 10.9734/BMRJ/2015/12941 |

| Source | Incubation | Infectious | Recovery | $R_0$ | β | Strain(s) | DOI |
|---|---|---|---|---|---|---|---|
| | | | | | | Karamzino 06/13, K 08/13, Vyazma 08/13, Stavropol 01/08 | |
| Walczak et al. (2020) | Yes | Yes | Yes | | | Pol18_28298_O111 | 10.3390/pathogens9030237 |
| Zani et al. (2018) | Yes | Yes | Yes | | | EstoniaPol18_28298_O111 | 10.1038/s41598-018-24740-1 |
| Zhao et al. (2019) | Yes | Yes | Yes | | | Pig/HLJ/18Estonia | 10.1080/22221751.2019.1590128 |

Model selection based on the F test resulted in the following final model being selected and analyzed for incubation period length:

$$\log(Incubation\ Period_{ij}) = 1 + Blood_{ij} + \text{Excreta}_{ij} + \text{Animal Age}_{ij} + \text{High Dose}_{ij} + (1|Study_j) + (1|Animal_{ij}|\ Study_j) + \epsilon_{ij}$$

Table S2. Log(Incubation) period coefficient estimates for fixed effects from the top LMM. The intercept represents temperature and low dose. Note the associated response is log(Incubation Period).

| Fixed Effects Term | Est | SE | t-Value | Degrees of Freedom | p-Value |
|---|---|---|---|---|---|
| (Intercept) | 1.70 | 0.06 | 26.41 | 29.41 | 0.00 |
| Animal Age (weeks) | 0.03 | 0.01 | 2.88 | 13.02 | 0.01 |
| Blood | -0.13 | 0.04 | -3.02 | 429.61 | 0.00 |
| Excreta | 0.26 | 0.04 | 6.23 | 470.44 | 0.00 |
| High Dose | -0.35 | 0.07 | -5.37 | 85.45 | 0.00 |
| **Random Effects Group** | **Parameter** | **SD** | | | |
| Study:Animal | Intercept | 0.11 | | | |
| Study | Intercept | 0.18 | | | |
| Residual | | 0.28 | | | |

Table S3. Model selection for random and fixed effects terms in the model for incubation period.

| Model | Eliminated | Number of Parameters | Log-Likelihood | AIC | Likelihood Ratio Test |
|---|---|---|---|---|---|
| | NA | 14 | -160.3 | 348.6 | NA |
| **(1 \| Source)** | 0 | 13 | -180.6 | 387.2 | 40.52 |
| **(1 \| Source:AnimalNumber)** | 0 | 13 | -166.2 | 358.5 | 11.86 |

| Model | Degrees of Freedom | Pr(>Chisq) |
|---|---|---|
| | NA | NA |
| **(1 \| Source)** | 1 | 1.941e-10 |
| **(1 \| Source:AnimalNumber)** | 1 | 0.0005731 |

| Model Term | Eliminated | Sum of Squares | Mean Squared Error | Degrees of Freedom | Den DF |
|---|---|---|---|---|---|
| **PCR** | 1 | 0.02336 | 0.02336 | 1 | 462.5 |
| **Recovery Rate** | 2 | 0.03056 | 0.03056 | 1 | 253.2 |
| **Isolation_Year** | 1 | 0.007366 | 0.007366 | 1 | 12.3 |
| **Recovery_Rate** | 2 | 0.02056 | 0.02056 | 1 | 252.3 |
| **PCR** | 3 | 0.04254 | 0.04254 | 1 | 457.5 |
| **Clinical_Signs** | 4 | 0.1236 | 0.1236 | 1 | 277.3 |
| **Intramuscular** | 5 | 0.1378 | 0.1378 | 1 | 207.1 |
| **Died** | 6 | 0.1993 | 0.1993 | 1 | 274.7 |
| **Animal_Age_weeks** | 0 | 0.6457 | 0.6457 | 1 | 13.02 |
| **Blood** | 0 | 0.7094 | 0.7094 | 1 | 429.6 |

| Model Term | F-value | Pr(>F) |
|---|---|---|
| **Isolation_Year** | 0.09552 | 0.7624 |
| **Recovery_Rate** | 0.2664 | 0.6062 |
| **PCR** | 0.5523 | 0.4578 |
| **Clinical_Signs** | 1.599 | 0.2071 |
| **Intramuscular** | 1.771 | 0.1847 |
| **Died** | 2.567 | 0.1103 |
| **Animal_Age_weeks** | 8.281 | 0.01294 |
| **Blood** | 9.097 | 0.002712 |
| **Excretions** | 38.77 | 1.058e-09 |
| **High_Dose** | 28.88 | 6.561e-07 |

Model selection based on the F-test resulted in the following final model being selected for animals that survived:

$$log(\text{Infectious } Period \text{ Survived}_{ij}) = 1 + Blood_{ij} + \text{PCR}_{ij} + (1|\text{Study}_j) + (1|Animal_{ij}|\text{ Study}_j) + \epsilon_{ij}$$

Table S4. Log(Infectious $\text{Period}_s$) Estimated fixed effects for the top infectious period model for animals that survived the studies. The intercept represents temperature and low dose.

| Term | t-Statistic | Degrees of Freedom | p-Value | Estimate | 95% CI |
|---|---|---|---|---|---|
| Intercept | 2.148 | 7.55 | 0.06599 | 0.78 | (-0.07, 1.62) |
| Blood | 5.899 | 115.4 | 3.72e-08 | 1.37 | (0.91, 1.83) |
| PCR | 6.312 | 115.2 | 5.298e-09 | 1.35 | (0.93, 1.77) |

Table S5. shows model selection for random and fixed effects terms in the model for infectious period for animals that survived.

| Model | Estimated | Number of Parameters | Log-Likelihood | AIC | Likelihood Ratio Test | Degrees of Freedom | Pr(>Chisq) |
|---|---|---|---|---|---|---|---|
| | NA | 13 | -183.6 | 392.2 | NA | NA | NA |
| **(1 \| Source)** | 0 | 12 | -186.1 | 396.3 | 5.102 | 1 | 0.0239 |
| **(1 \| Source:AnimalNumber)** | 0 | 12 | -184 | 392.1 | 6.174e-09 | 1 | 0.9999 |

| Term | Eliminated | Sum of Squares | Mean Squared Error | Degrees of Freedom | DenDF | F-Value | Pr(>F) |
|---|---|---|---|---|---|---|---|
| **Animal_Age_weeks** | 1 | 0.004393 | 0.004393 | 1 | 2.186 | 0.00398 | 0.955 |
| **Recent_Isolation** | 2 | 0.001449 | 0.001449 | 1 | 3.22 | 0.001312 | 0.9732 |
| **High_Dose** | 3 | 0.1277 | 0.1277 | 1 | 60.07 | 0.1157 | 0.735 |
| **Recovery_Rate** | 4 | 0.6025 | 0.6025 | 1 | 27 | 0.5478 | 0.4656 |
| **Direct_Contact** | 5 | 0.7819 | 0.7819 | 1 | 23.03 | 0.7186 | 0.4053 |
| **Isolation_Year** | 6 | 0.7747 | 0.7747 | 1 | 4.418 | 0.7178 | 0.4403 |
| **Intramuscular** | 7 | 2.686 | 2.686 | 1 | 113.3 | 2.493 | 0.1172 |
| **Blood** | 0 | 38 | 38 | 1 | 115.4 | 34.8 | 3.72e-08 |
| **PCR** | 0 | 43.51 | 43.51 | 1 | 115.2 | 39.85 | 5.298e-09 |

### Table S6. Strain specific mortality rate

| ASFV Strain | Isolation Year | Viral.Strain | Articles | Animals | Percent Recovering |
|---|---|---|---|---|---|
| Armenia | 2008 | Yerevan Armenia 2029 | 1 | 36 | 0.0 |
| Armenia | 2008 | Yerevan Armenia 2030 | 1 | 36 | 0.0 |
| Armenia | 2008 | Yerevan Armenia 2031 | 1 | 36 | 0.0 |
| Belgium 2018/1 | 2018 | Belgium 2018 | 1 | 89 | 50.0 |
| China | 2018 | Pig/HLJ/18 | 1 | 24 | 0.0 |
| China | 2020 | HLJ HRB1 20 | 1 | 24 | 62.5 |
| Dominican Republic 2021 | 2021 | Dominican Republic strain | 1 | 48 | 30.0 |
| Estonia | 2014 | Estonia | 1 | 194 | 63.6 |
| Estonia | 2014 | Ida-Viru Estonia | 1 | 100 | 10.0 |
| Estonia | 2015 | Es15/WB-Tartu-14 | 1 | 20 | 33.3 |
| Estonia | 2015 | Es15/WB-Valga-6 | 1 | 20 | 33.3 |
| Estonia | 2016 | Est16/WB/Viru8 | 1 | 20 | 33.3 |
| Georgia 2007 | 2007 | Georgia 2007 | 7 | 1001 | 0.0 |

| | | | | | |
|---|---|---|---|---|---|
| Latvia | 2017 | Lv17/WB/Rie1 | 1 | 20 | 100.0 |
| Lithuania | 2014 | Lithuania LT14/1490 | 1 | 164 | 11.1 |
| Poland | 2015 | POL/2015/Podlaskie/Lindholm | 1 | 260 | 20.6 |
| Poland | 2016 | Pol16/DP/OUT21 | 1 | 20 | 0.0 |
| Poland | 2018 | Pol18 28298 O111 | 1 | 164 | 9.1 |
| Russian Federation | 2008 | Stavropol 01/08 | 1 | 12 | 0.0 |
| Russian Federation | 2013 | Boguchary 06/13 | 1 | 12 | 0.0 |
| Russian Federation | 2013 | K 08/13 | 1 | 4 | 0.0 |
| Russian Federation | 2013 | Karamzino 06/13 | 1 | 12 | 0.0 |
| Russian Federation | 2013 | Kashino 04/13 | 1 | 8 | 0.0 |
| Russian Federation | 2013 | Vyazma 08/13 | 1 | 12 | 0.0 |
| Russian Federation | 2013 | Zubtsovo 06/13 | 1 | 113 | 0.0 |
| Russian Federation | 2014 | Grafsky 06/14 | 1 | 128 | 0.0 |
| Russian Federation | 2014 | Odintsovo 02/14 | 2 | 114 | 8.0 |
| Russian Federation | 2014 | Voronezh-Agro 12/14 | 1 | 108 | 0.0 |
| Russian Federation | 2015 | Krasnodar 07/15 | 1 | 128 | 0.0 |
| Russian Federation | 2015 | Lysogorye 07/15 | 1 | 128 | 6.2 |
| Russian Federation | 2015 | Ryazan 10/15 | 1 | 128 | 0.0 |
| Russian Federation | 2015 | Sobinka 07/15 | 1 | 128 | 0.0 |
| Russian Federation | 2016 | Bolokhovsky 07/15 | 1 | 64 | 25.0 |
| Russian Federation | 2016 | Lipetsk 12/16 | 1 | 64 | 50.0 |
| Russian Federation | 2016 | Martins-Krym 01/16 | 1 | 36 | 0.0 |
| Russian Federation | 2016 | Ryazan 03/16 | 1 | 64 | 0.0 |
| Russian Federation | 2016 | Ryazan 07/16 | 1 | 64 | 0.0 |
| Russian Federation | 2017 | Kaliningrad 10/17 | 1 | 36 | 0.0 |
| Russian Federation | 2018 | Timashevsk 01/18 | 1 | 100 | 10.0 |
| Vietnam | 2019 | VNUA/HY/Vietnam | 1 | 100 | 0.0 |

Table S7 Fixed effects terms for the Generalized Linear Mixed Model fit to mortality rate.

| Fixed Effects Term | t-Statistic | P-Value | Estimate | 95% CI |
|---|---|---|---|---|
| **Intercept** | -4.09 | 4.343e-05 | -2.99 | (-4.43, -1.56) |
| **High Dose** | 3.15 | 0.001635 | 2.00 | (0.76, 3.24) |
| **Intramuscular** | 1.53 | 0.1266 | 1.52 | (-0.43, 3.48) |
| **Direct Contact** | 3.83 | 0.0001303 | 2.77 | (1.35, 4.18) |
| **Animal Age** | 2.43 | 0.01489 | 0.49 | (0.1, 0.89) |
| **High Dose*Intramuscular** | -3.21 | 0.0156 | -3.21 | (-5.8, -0.61) |

Table S8. Mortality rate model selection.

| Intercept | Animal Age | Direct Contact | High Dose | Intramuscular | High Dose * Intramuscula | df | Log-Likelihoo | AICc | Δ AIC c | Weight |
|---|---|---|---|---|---|---|---|---|---|---|
| -2.993 | 0.4912 | Yes | Yes | Yes | Yes | 7 | -131.3 | 277 | 0 | 0.5565 |
| -2.671 | 0.568 | Yes | Yes | | | 5 | -134.5 | 279.1 | 2.156 | 0.1893 |
| -2.469 | 0.5221 | Yes | Yes | Yes | | 6 | -134.2 | 280.7 | 3.768 | 0.0846 |
| -2.666 | | Yes | Yes | Yes | Yes | 6 | -134.3 | 280.9 | 3.946 | 0.07737 |
| -1.788 | 0.68 | Yes | | | | 4 | -137.1 | 282.3 | 5.314 | 0.03904 |
| -1.489 | 0.599 | Yes | | Yes | | 5 | -136.6 | 283.4 | 6.439 | 0.02225 |
| -2.134 | | Yes | Yes | Yes | | 5 | -137.3 | 284.9 | 7.904 | 0.0107 |
| -2.637 | | Yes | Yes | | | 4 | -138.6 | 285.4 | 8.428 | 0.00822 |

Table S9. Log(Beta) fixed effects model terms.

| Term | Estimate | SE | z-Value | p-Value | 95% CI |
|---|---|---|---|---|---|
| **Intercept** | -0.271 | 0.109 | -22.485 | 0.0129 | (-0.484, -0.57) |
| **Isolation Year** | -0.513 | 0.0611 | -8.39 | <.0001 | -0.63,-0.39) |
| **Latent Period** | 0.446 | 0.1341 | 3.32 | 0.0009 | (0.1835, 0.7090) |
| **Between Pen** | -0.777 | 0.15 | -5.24 | <.0001 | (-1.066, -0.486) |

Table S10. Log(Beta) model selection.

| Intercept | Between Pen | Isolation Year | Latent Period | Within Pen | Degrees of Freedom | Log-Likelihood | AICc | Δ AICc | Weight |
|---|---|---|---|---|---|---|---|---|---|
| Yes | | -0.513 | 0.446 | NA | 5 | -11.16611 | 35.05949 | 0.000000 | 0.589 |
| Yes | Yes | -0.268 | 0.646 | NA | 5 | -12.05875 | 36.84477 | 1.785276 | 0.241 |
| Yes | Yes | -0.48 | 0.473 | NA | 6 | -11.14329 | 38.28659 | 3.227091 | 0.117 |
| Yes | Yes | -0.715 | | NA | 5 | -14.29443 | 41.31613 | 6.256639 | 0.026 |
| Yes | | -0.449 | | NA | 4 | -16.39869 | 42.53651 | 7.477014 | 0.014 |
| Yes | Yes | | 0.783 | NA | 5 | -15.05810 | 42.84347 | 7.783977 | 0.012 |
| Yes | Yes | | 0.543 | NA | 4 | -19.21593 | 48.17099 | 13.111499 | 0.001 |
| Yes | | -0.384 | 0.576 | NA | 4 | -20.84933 | 51.43778 | 16.378286 | 0.000 |
| Yes | Yes | -0.173 | | NA | 4 | -22.66194 | 55.06302 | 20.003521 | 0.000 |

Table S11. Parameters for Gamma distribution fits.

| Parameter | Animal Status | Measurement Method | Sample Type | Shape | Rate | Median | 2.5 and 97.5% Quantiles | KS Test p-Value |
|---|---|---|---|---|---|---|---|---|
| **Latent Period** | All | All | All | 8.01 ± 0.47 | 1.70±0.10 | 4.52 | (2.04, 8.5) | 0.28 |
| **Infectious Period** | Died | All | All | 2.69 ± 0.17 | 0.53 ± 0.04 | 4.5 | (0.93, 12.73) | 0.07 |
| **Infectious Period** | Survived | PCR | Blood | 1.37 ±0.31 | 0.034 ±0.01 | 31.2 | (2.4, 130.79) | 0.26 |
| **Infectious Period** | Survived | Viral Isolation | Blood | 1.95±0.49 | 0.087±0.025 | 18.76 | (2.62, 63.11) | 0.02 |
| **Infectious Period** | Survived | PCR | Excreta | 0.74±0.14 | 0.0234±0.01 | 19.94 | (0.31, 135.56) | 0.57 |
| **Infectious Period** | Survived | Viral isolation | Excreta | 1.21±0.34 | 0.369±0.13 | 2.44 | (0.14, 11.18) | 0.15 |
